# Enteroendocrine cells couple nutrient sensing to nutrient absorption by regulating ion transport

**DOI:** 10.1101/2020.04.21.053363

**Authors:** Heather A. McCauley, Andrea L. Matthis, Jacob R. Enriquez, Jonah Nichol, J. Guillermo Sanchez, William J. Stone, Nambirajan Sundaram, Michael A. Helmrath, Marshall H. Montrose, Eitaro Aihara, James M. Wells

## Abstract

The ability to absorb ingested nutrients is an essential function of all metazoans and utilizes a wide array of nutrient transporters found on the absorptive enterocytes of the small intestine. A unique population of patients has previously been identified with severe congenital malabsorptive diarrhea upon ingestion of any enteral nutrition. The intestines of these patients are macroscopically normal, but lack enteroendocrine cells (EECs), suggesting an essential role for this rare population of nutrient-sensing cells in regulating macronutrient absorption. We used human and mouse models of EEC deficiency to identify a new role for the EEC hormone peptide YY in regulating ion-coupled absorption of glucose and dipeptides the small intestine. We found that peptide YY is required in to maintain normal electrophysiology in the presence of vasoactive intestinal polypeptide, a potent stimulator of ion secretion produced by enteric neurons. Administration of peptide YY to EEC-deficient mice restored normal electrophysiology, improved glucose and peptide absorption, diminished diarrhea and rescued postnatal survival. These data suggest that peptide YY is a key regulator of macronutrient absorption in the small intestine and may be a viable therapeutic option to treat patients with malabsorption.

## Introduction

Enteroendocrine cells (EECs) are a rare population of cells found in the gastrointestinal epithelium that sense nutrients that are passing through the gut and in response secrete more than 20 distinct biologically active peptides. These peptides act in an endocrine or paracrine fashion to regulate all aspects of nutrient homeostasis including satiety, mechanical and chemical digestion, nutrient absorption, storage and utilization (Gribble and Reimann, 2019). Humans (Wang et al., 2006) and mice (Mellitzer et al., 2010) with genetic mutations that impact formation or function of EECs have intractable malabsorptive diarrhea, metabolic acidosis, and require parenteral nutrition or small-bowel transplant for survival. These findings were the first to link EECs to the absorption of macronutrients; however, the mechanism by which EECs contribute to this vital process is unknown. Poor absorption of macronutrients is a global health concern, with underlying etiology including short-gut syndrome, enteric pathogen infection, and malnutrition. Therefore, identification of factors regulating nutrient absorption has significant therapeutic potential.

Absorption of carbohydrate and protein requires coordinated activity of nutrient and ion transporters in the small intestine. Glucose is primarily absorbed via sodium-glucose cotransporter SGLT1, which uses a downhill Na^+^ gradient to transport one glucose or galactose molecule with two sodium ions from the lumen into the enterocyte (Wright et al., 2011). The majority of dietary protein absorption occurs via Na^+^- and H^+^-linked amino acid transporters and PEPT1, which imports di- and tri-peptides coupled with a hydrogen ion (Chen et al., 2010). The electrochemical gradients that drive nutrient absorption are maintained in part by ion transporters, including the cystic fibrosis transmembrane receptor (CFTR), which exports chloride (Wright et al., 1997), and sodium-hydrogen exchanger NHE3, which maintains Na^+^ and H^+^ microclimates across the apical membrane (Thwaites et al., 2002). Activity of CFTR and NHE3 are, in turn, regulated by levels of cyclic AMP (cAMP) (Burleigh and Banks, 2007; Yun et al., 1997).

Most secreted EEC peptides signal via G protein-coupled receptors that act via second messenger cascade effectors like cAMP. Given the requirement for EECs in regulating macronutrient absorption, we investigated the possibility that EEC-derived peptides coupled nutrient sensing to nutrient absorption by regulating electrogenic transport in neighboring enterocytes. Two well-studied peptides governing ion and water homeostasis in the colon are vasoactive intestinal peptide (VIP) and peptide YY (PYY). VIP, secreted from enteric neurons, signals via the G_αs_-coupled VIPR1 (VPAC1) on epithelial cells to raise levels of intracellular cAMP. In contrast, EEC-derived PYY acts in a paracrine fashion on colonocytes to lower cAMP via the epithelial G_αi_ coupled receptor NPY1R (Cox et al., 2010; Hyland et al., 2003; Moodaley et al., 2017; Tough et al., 2011). We posited that the mechanism underlying malabsorptive diarrhea in patients lacking EECs might be due to loss of a similar EEC-ENS regulatory feedback in the small intestine, thus disrupting electrogenic nutrient absorption. Here, we found that PYY regulates normal ion transport and ion-coupled nutrient absorption in mouse and human small intestine, and that administration of exogenous PYY was sufficient to restore normal electrophysiology, nutrient absorption, and survival in EEC-deficient animals.

## Results

### The PYY-VIP axis regulates ion and water transport in mouse and human small intestine

If EECs were required for regulating the normal electrophysiology of the small intestine, we would expect to see deranged ion transport in intestinal tissues lacking EECs. To investigate this, we used EEC-deficient mice (*VillinCre;Neurog3^flox/flox^*) (Mellitzer et al., 2010) and three different human small intestinal tissue models all derived from pluripotent stem cells (PSCs): human intestinal organoids (HIOs) derived *in vitro* (Spence et al., 2011), HIOs that were matured to robust crypt-villus architecture *in vivo* (Watson et al., 2014), and epithelial organoids (enteroids) derived from crypts of matured HIO tissues (Watson et al., 2014). We generated EEC-deficient human small intestinal tissue by using PSC lines that had a null mutation in *NEUROG3* (McGrath et al., 2015), the basic helix-loop-helix transcription factor required for EEC formation in mice (Jenny et al., 2002) and humans (Wang et al., 2006). As previously reported (Zhang et al., 2019), NEUROG3^−/−^ small intestinal organoids completely lacked EECs, but were otherwise normal in appearance (Figure S1).

Ion and water transport in the colon are regulated by EEC-derived PYY and ENS-derived VIP. To formally test whether the PYY-VIP axis operated in human and mouse small intestine, we performed experiments in EEC-deficient tissues without a functional ENS wherein we controlled PYY and VIP levels experimentally. We first determined the effects of the PYY-VIP axis on small intestine by measuring CFTR-mediated ion and water efflux (Dekkers et al., 2013) following exposure of human HIO-derived enteroids to the potent secretagogue VIP (Figure 1A). EEC-deficient enteroids swelled significantly more than did wild-type but blocking NPY1R in wild-type enteroids mimicked the EEC-deficient response (Figure 1A). Exogenous PYY blocked VIP-induced swelling in both wild-type and EEC-deficient enteroids in an NPY1R-dependent manner (Figure 1A), demonstrating that the PYY-VIP axis regulates ion and water secretion in human small intestine. We next tested the activity of NHE3 as a measure of Na^+^-dependent intracellular pH recovery after acidic challenge (Foulke-Abel et al., 2016) and found that EEC-deficient enteroids displayed impaired NHE3 function (Figure 1B). There was no difference in expression of *CFTR, SLC9A3* (encoding NHE3), *VIPR1* or *NPY1R* between wild-type and EEC-deficient human small intestinal organoids or enteroids (Figure 1C and Figure S2A-B). Together, these data suggest that PYY plays an important role in the regulation of ion transport in the small intestine, and that the abnormal response to VIP in EEC-deficient enteroids can be normalized by the addition of exogenous PYY.

**Figure 1.**
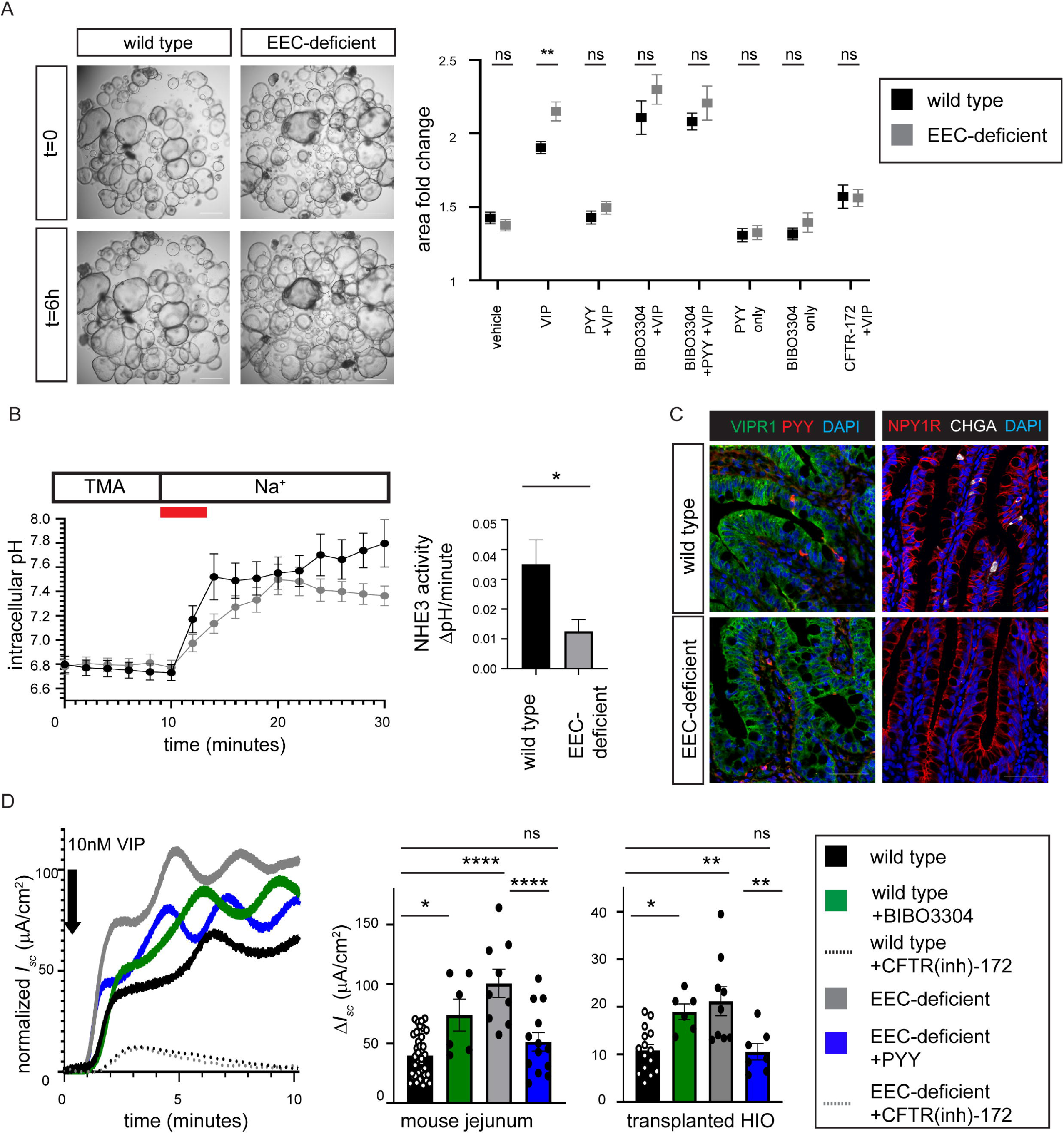
The PYY-VIP axis regulates ion and water transport in mouse and human small intestine. A. PYY and VIP regulate ion and water transport in HIO-derived small intestinal enteroids. Addition of VIP to enteroids induced ion and water transport as measured by swelling. EEC-deficient enteroids had an elevated response to VIP compared to wild-type enteroids (**P=0.004). Upon addition of PYY, there was no difference in swelling between wild-type and EEC-deficient enteroids, and inhibition of VIP-induced swelling. Chemical inhibition of the PYY receptor NPY1R with BIBO3304 resulted in swelling of wild-type enteroids to EEC-deficient levels, and abolished the inhibitory effects of PYY in both wild-type and EEC-deficient enteroids. VIP-induced enteroid swelling was CFTR dependent and blocked by the CFTR inhibitor CFTR-172. Scale bars = 500 μm. Black bars represent wild-type and gray bars represent EEC-deficient enteroids. Error bars are ± SEM. n=283 wild-type and n=351 EEC-deficient enteroids over three biologically independent enteroid lines. Statistics calculated by two-way ANOVA with Sidak’s multiple comparisons test. B. EEC-deficient enteroids displayed impaired NHE3 activity. EEC-deficient enteroids exhibited reduced Na^+^-dependent recovery of intracellular pH after an acid load using the ratiometric pH indicator SNARF-4F. Quantification is of initial rate of Na^+^-dependent pH recovery (red line). n=16 wild-type, n=18 mutant enteroids; *P=0.01. Error bars are ± SEM; statistics calculated by unpaired, two-tailed Student’s *t*-test. C. The localization of the VIP receptor VIPR1 and PYY receptor NPY1R was comparable between wild-type and EEC-deficient human intestinal epithelium. PYY+ and CHGA+ cells were only found in wild-type HIOs. Scale bars = 100 μm. D. PYY modulates the stimulatory effects of VIP in mouse and human small intestine. In the Ussing chamber, EEC-deficient small intestine displayed a greater response (Δ*I*_sc_) to 10 nM VIP than did wild-type (mouse, n=20 wild-type, 8 mutant, ****P<0.0001; human, n=15 wild-type, 9 mutant, **P=0.001). Inhibition of NPY1R in wild-type tissue with BIBO3304 resulted in an elevated response to VIP compared to untreated wild-type (mouse, n=24, *P=0.01; human, n=7, *P=0.04). Addition of exogenous PYY reduced the magnitude of EEC-deficient response to VIP (n=8 mutant mice, ****P<0.0001; n=7 mutant HIOs, **P=.007) to wild-type levels. Electrogenic responses to VIP were blocked by the CFTR inhibitor CFTR-172m (dotten lines). One representative trace is shown (mouse), with baseline *I*_sc_ normalized to 0 μA/cm^2^. Error bars are ± SEM; statistics calculated by one-way ANOVA with Tukey’s multiple comparisons test.

If PYY were required to regulate electrochemical transport in the small intestine, we would expect that disruption of PYY signaling in wild-type small intestinal tissue would cause abnormal basal short-circuit current (*I*_sc_). To investigate this we isolated full thickness intestinal mucosa from *in vivo* matured human intestinal organoids and from the jejunum of wild-type mice and measured basal *I*_sc_ in a modified Ussing chamber (Clarke, 2009). Chemical inhibition of NPY1R in wild-type mouse jejunum and human intestinal organoids was sufficient to elevate the basal *I_sc_* to EEC-deficient levels (Figure S2C). Conversely, treatment of EEC-deficient mouse and human tissues with exogenous PYY reduced the basal *I_sc_* to wild-type levels in an NPY1R-dependent manner (Figure S2C). These data indicated that endogenous PYY signaling plays an essential role in maintaining normal electrophysiology in the small intestine.

We then investigated if PYY was required to modulate the stimulatory effects of VIP in mouse and human small intestine. We inhibited voltage-gated neuronal firing in mouse jejunum by including tetrodotoxin (Hyland et al., 2003) in all experiments so that we could precisely monitor epithelial response to exogenous VIP. Chemical inhibition of NPY1R in isolated wild-type tissues was sufficient to cause an elevated response to VIP (Figure 1D). This indicated that endogenous PYY signaling was required in the small intestine to modulate the stimulatory effects of VIP. Consistent with this, EEC-deficient mouse and human small intestinal tissue similarly displayed an exaggerated *I*_sc_ response to exogenous VIP compared to wild-type (Figure 1D). Addition of exogenous PYY to EEC-deficient small intestine was sufficient to restore the *I*_sc_ to normal (Figure 1D). These data suggested that PYY is required for maintaining a normal electrochemical response to VIP in the small intestine and that PYY can normalize this process in EEC-deficient small intestinal tissue. Furthermore, these data suggest that imbalance of this axis may be a mechanism underlying malabsorptive diarrhea suffered by patients without EECs.

### PYY restores normal glucose absorption in EEC-deficient human and mouse small intestine

While it is known that EECs sense nutrients, the mechanism linking sensing to the control of nutrient absorption is unclear. A hint came from the effects of enteral feeding of EEC-deficient patients, which resulted in a massive diarrheal response. This suggests that an inability to sense luminal nutrients uncoupled the ability to adequately absorb them. To explore this possibility we evaluated ion-coupled nutrient absorption in EEC-deficient small intestine. We observed an accelerated initial response to luminal glucose in the presence of VIP in EEC-deficient mouse and human intestinal tissues in the Ussing chamber (Figure 2C), as predicted if the normal electrochemical gradients were perturbed (Figure 2A-B). This recapitulated the exacerbated diarrhea observed in patients without EECs when they were fed with carbohydrate (Wang et al., 2006). Exogenous PYY restored a normal glucose response in EEC-deficient mouse and human tissue, and inhibition of NPY1R in wild-type caused an exaggerated initial response to glucose (Figure 2C). These data indicate that PYY is both necessary and sufficient to modulate glucose absorption in the small intestine. We found no defects in expression of SGLT1, GLUT2, (Figure 2D and S2A) or maximum absorptive competency of Na^+^-coupled glucose transport (Figure 2E-G) in human epithelium without EECs. These data suggest that SGLT1 is competent to absorb glucose, but activity is dysregulated in the context of abnormal ion transport in the absence of EECs.

**Figure 2.**
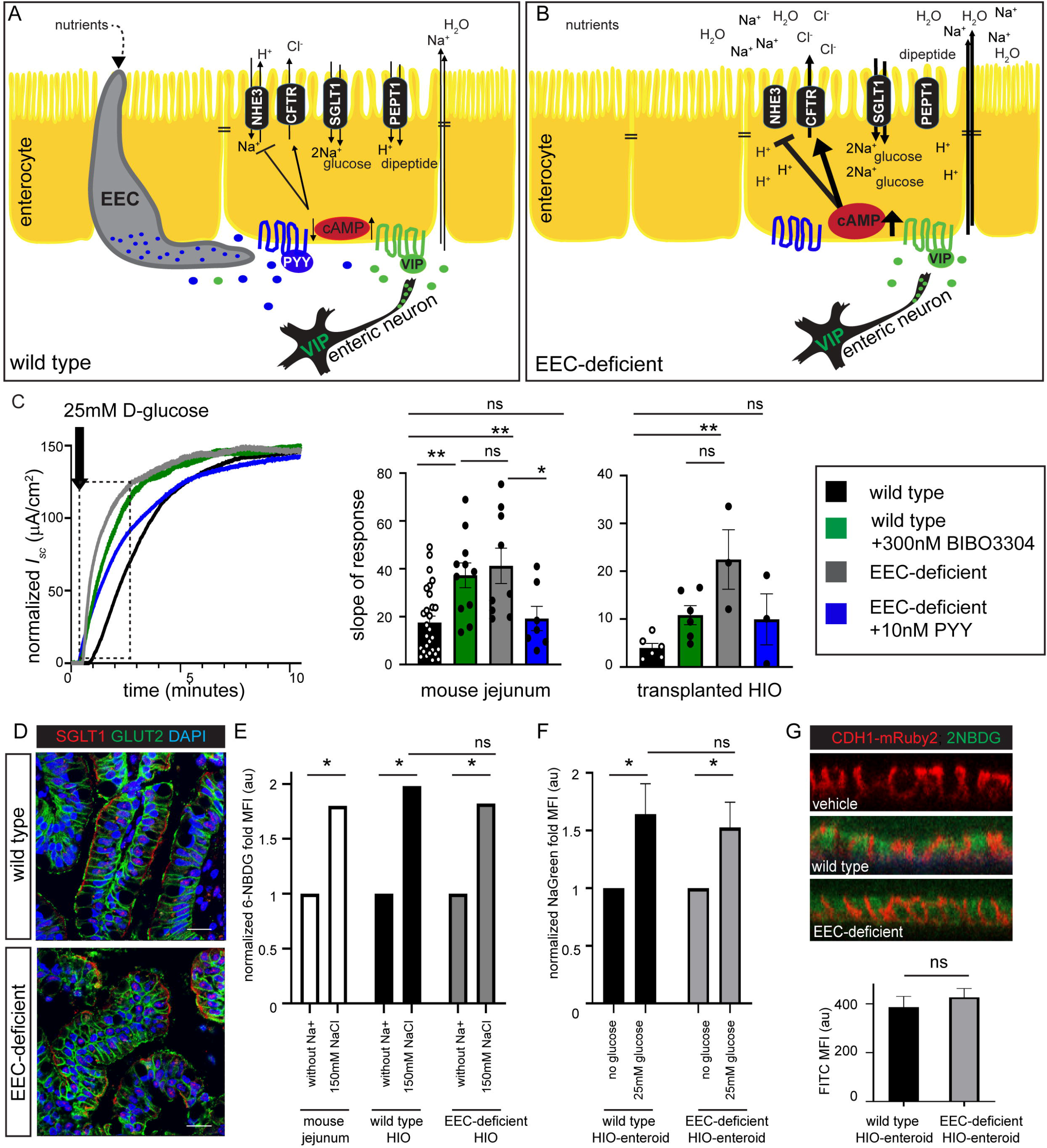
PYY restores normal glucose absorption in EEC-deficient human and mouse small intestine. A. Schematic depicting the PYY-VIP paracrine axis regulating ion and water homeostasis. EEC-derived PYY and ENS-derived VIP both act via G-protein coupled receptors (NPY1R and VIPR1, respectively) on enterocytes. VIP signaling raises intracellular cAMP levels resulting in activation of CFTR and efflux of chloride ions while concurrently inhibiting the sodium-hydrogen exchanger NHE3. The downstream results are that water and sodium are drawn to the intestinal lumen via paracellular spaces to balance the secreted chloride. PYY is secreted in response to luminal nutrients and acts as a counterbalance to VIP by lowering intracellular cAMP levels. Transport of luminal nutrients into the enterocyte depends on these ion gradients; SGLT1 transports glucose with two Na^+^ ions and PEPT1 transports di-/tri-peptides with an H^+^ ion. B. In the absence of EECs, ion and water homeostasis is deregulated due to loss of one arm of the PYY-VIP axis. In EEC-deficient small intestine, loss of PYY results in increased cAMP-signaling, increased chloride transport, and increased water and sodium accumulation in the intestinal lumen. Reduced NHE3 transport activity would cause accumulation of cytosolic H^+^ and a decrease in pH. Subsequently, nutrient absorption would be dysregulated, with diminished di-/tri-peptide absorption due to increased intracellular proton accumulation and with increased uptake of glucose due to an exaggerated Na^+^ gradient across the apical membrane. C. Na+-coupled glucose transport is deranged in EEC-deficient human and mouse small intestine. Wild-type and EEC-deficient human and mouse intestinal tissues were treated with VIP, then 25 mM D-Glucose was added to the luminal chamber. EEC-deficient intestine had an elevated initial response to glucose (mouse, n= 28 wild-type, n=9 mutant, **P=0.001; HIO, n=6 wild-type, n=4 mutant, **P=0.002) that was returned to wild-type levels by pre-treatment with 10 nM exogenous PYY (mouse, n=7, *P=0.04; HIO, n=3). Inhibition of the NPY1R in wild-type tissues using the antagonist BIBO3304 caused an abnormal initial response to glucose that mimicked EEC-deficient tissues (mouse, n=12, **P=0.005; HIO, n=6). Bar graphs represent the slope of the curve depicted within the boxed area. Error bars are ± SEM; statistics calculated by one-way ANOVA with Tukey’s multiple comparisons test. D. The subcellular distribution of glucose transporters SGLT1 and GLUT2 is normal in human intestinal tissue lacking EECs. Scale bars = 50 μm. E. SGLT1 is functional in EEC-deficient human small intestine. Human small intestinal tissue was isolated and transport of glucose in response to saturating amounts of NaCl were measured using the glucose analog 6-NBDG. EEC-deficient human small intestinal cells displayed similar total 6-NBDG uptake in the presence of NaCl (*P=0.01) to wild-type human intestinal cells (*P=0.01) and wild-type mouse jejunum cells (*P=0.01), demonstrating functional SGLT1-mediated transport. Statistics calculated by one-way ANOVA with Tukey’s multiple comparisons test. F. The ability of SGLT1 to transport Na^+^ is not altered in EEC-deficient enteroids. Enteroids were stained with the Na^+^ fluorescent indicator NaGreen in the presence or absence of 25 mM glucose. The Na^+^ transport activity of SGLT1 in the presence of glucose is similar in both wild-type and EEC-deficient epithelium as measured by fluorescence intensity (MFI) (*P=0.01). Data represents 4 independent experiments. Statistics calculated by one-way ANOVA with Tukey’s multiple comparisons test. G. Total glucose transport is similar in wild-type and EEC-deficient monolayer cultures. Wild-type and EEC-deficient enteroids were cultured as monolayers on transwell inserts and exposed to 25 mM D-glucose with 1 mM fluorescent glucose analog 2-NBDG on the apical surface. The fluorescence intensity of the basal chamber was quantified after 30 minutes (lower graph). The epithelium was then analyzed for 2-NBDG within CDH1-mRuby2-positive epithelium. Data represents 8 independent experiments. Statistics calculated by unpaired t-test.

### H^+^-coupled dipeptide absorption is impaired in EEC-deficient small intestine

Approximately 80% of ingested amino acids were recovered in the stool of the index EEC-deficient patient (Wang et al., 2006), suggesting a critical role for EECs in regulating protein absorption. Consistent with this, we observed a striking loss of ion-coupled dipeptide absorption when human and mouse EEC-deficient small intestine were challenged with VIP (Figure 3A), despite normal expression of PEPT1 (Figures 3B and S2A). VIP has an established role in inhibition of NHE3 and PEPT1-mediated dipeptide absorption (Anderson et al., 2003; Thwaites et al., 2002), but we were surprised to find that EEC-deficient intestine remained unable to respond to dipeptide when PYY was provided (Figure 3A). This suggested that dysregulated H^+^ gradients may be a more stable phenotype in EEC-deficient intestine, and not easily reversed by PYY within minutes. To explore this possibility, we treated enteroids with or without PYY for one week *in vitro* in the presence of VIP. Wild-type enteroids were able to maintain their intracellular pH in the presence of VIP but EEC-deficient enteroids became significantly more acidic (Figure 3C). However, EEC-deficient enteroids were restored to normal intracellular pH levels and normal *SLC9A3* expression (encoding NHE3) in the presence of PYY (Figures 3C and S3). This suggested that long-term exposure to an imbalanced EEC-ENS axis dysregulates intestinal physiology, and that, over time, PYY may be sufficient to restore intracellular pH and dipeptide absorption in EEC-deficient small intestine.

**Figure 3.**
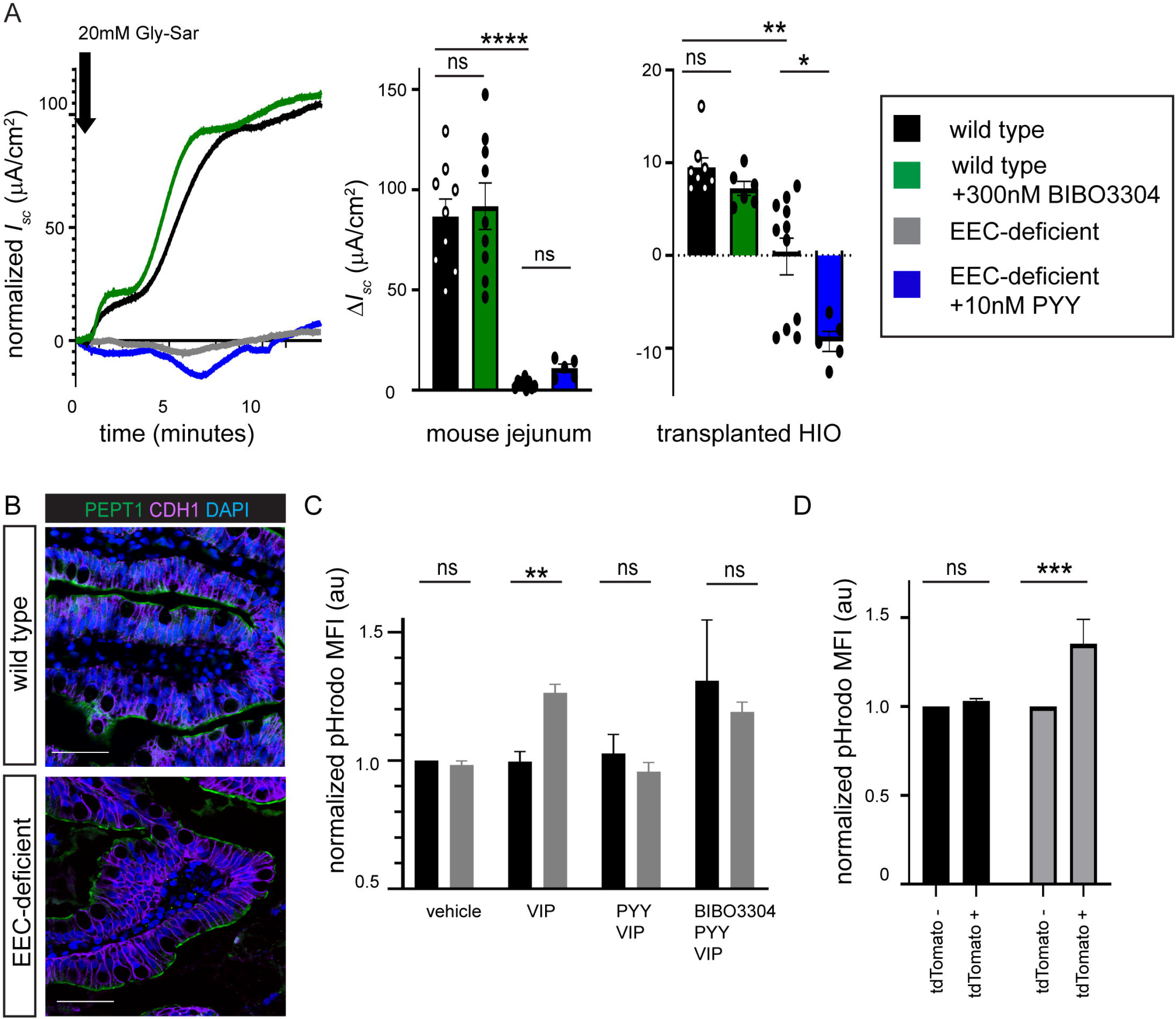
H^+^-coupled dipeptide absorption is impaired in EEC-deficient small intestine. A. EEC-deficient human and mouse small intestine did not respond to luminal Gly-Sar, a nonhydrolyzable dipeptide, in the Ussing chamber when exposed to 10nM VIP (mouse, n= 9 wild-type, n=6 mutant, ****P<0.0001; human, n=11 wild-type, n=5 mutant, **P=0.006). 10 minutes pre-treatment of EEC-deficient tissue with 10 nM exogenous PYY (mouse, n=6), or of wild-type tissue with 300 nM NPY1R inhibitor BIBO3304 (mouse, n=9; human, n=6) did not alter the *I*_sc_ response to Gly-Sar. Error bars are ± SEM; statistics calculated by one-way ANOVA with Tukey’s multiple comparisons test. B. Expression and localization of peptide transporter PEPT1 is unchanged in EEC-deficient human small intestine. Scale bars = 50 μm. C. The PYY-VIP axis regulates intracellular pH in human small intestinal cells. Wild-type and EEC-deficient enteroids were differentiated in the presence of 10 nM VIP for 5-7 days. EEC-deficient enteroids treated with VIP developed an H^+^ imbalance with an acidic cytoplasm whereas wild-type enteroids were able to maintain their intracellular pH (**P=0.004). Concurrent treatment with 10 nM PYY normalized the pH in EEC-deficient enteroids and was dependent on NPY1R. pHrodo mean fluorescence intensity (MFI) was normalized to vehicle-treated wild-type. n= 3 independent experiments. Error bars are ± SEM; statistics calculated by the Holm-Sidak method. D. Small intestinal EECs regulate proton transport in a paracrine fashion. Using animals with mosaic loss of EECs we found that regions of epithelium that escaped recombination had normal pH and H^+^ transport. Adjacent regions that expressed tdTomato, indicating Cre activity, had impaired elevated cytosolic H^+^ as measured by flow cytometry using the fluorescent pH indicator dye pHrodo. There was no difference in pHrodo MFI between mosaic regions in wild-type jejunum (n=8), but a significant increase in pHrodo MFI, indicating relative acidic pH, in EEC-deficient jejunum compared to non-recombined epithelial cells within the same segment of jejunum (n=4, ***P=0.0002). Error bars are ± SEM; statistics calculated by two-way ANOVA with Sidak’s multiple comparisons test.

We have demonstrated that inhibiting PYY signaling in isolated wild-type small intestinal tissues was sufficient to perturb normal electrophysiology in both human and mouse. This suggests that *in vivo* the mechanism of action of PYY could be paracrine rather than endocrine. PYY-expressing EECs are abundant in mouse and human small intestine (Egerod et al., 2012) (Figure S4). Moreover, PYY-expressing EECs extend long basal processes which underlie several neighboring epithelial cells (Bohorquez et al., 2014; Bohorquez et al., 2015), raising the possibility that they may exert paracrine effects on whole populations of nearby enterocytes. We therefore investigated whether the effects of PYY on ion transport in the small intestine occurred via paracrine mechanisms. To do this, we exploited the mosaicism of *VillinCre* mice to determine if regions of EEC-deficient epithelium had different transporter activities as compared to regions of epithelium that still had EECs. We observed in *VillinCre;Neurog3^flox/flox^* mice that 4.38±2.56% of jejunum escaped tdTomato labeling (Figure S5) and that in regions that had EECs, neighboring enterocytes had a normal intracellular pH indicating normal ion transport. In contrast, enterocytes in EEC-deficient regions were significantly more acidic indicating perturbed H^+^ transport (Figures 3D and S5). Together these data suggest that EECs control local H^+^ transporter activity and dipeptide responsiveness in the small intestine via paracrine mechanisms.

### Exogenous PYY rescues EEC-deficient mice from malabsorptive diarrhea and death and restores normal glucose and dipeptide transport

As previously reported (Mellitzer et al., 2010), *VillinCre;Neurog3^flox/flox^* mice suffer from malabsorptive diarrhea and exhibit severely impaired postnatal survival, with only a small fraction of mice surviving weaning. Our data suggested that treatment with PYY might restore normal carbohydrate and protein absorption the intestines of EEC-deficient animals. We therefore used *VillinCre;Neurog3^flox/flox^* mice as a preclinical model to test if PYY could reverse malabsorptive diarrhea and improve postnatal survival (Figure 4A-B). We began daily treatment of mutant mice at postnatal day 10 with 10 μg PYY(1-36) by intraperitoneal injection. PYY can be converted to PYY(3-36) by the protease DPP4 (Mentlein et al., 1993), and this form of PYY has potent anorexic effects in the brain (Batterham et al.). We therefore co-injected PYY(1-36) and a DPP4 inhibitor to prevent PYY cleavage and to better target the epithelial NPY1R receptor that preferentially binds the 1-36 form (Hyland et al., 2003; Mentlein et al., 1993; Tough et al., 2011). Patients with EEC-deficiency die without total parenteral nutrition, and similarly very few EEC-deficient mice survive without treatment within the first few weeks. However, PYY injections dramatically improved mutant survival up to 88% (Figure 4A). Moreover, PYY treatment reduced diarrhea and improved fecal output of mutant mice to either be indistinguishable from wild-type or only slightly wet but well-defined pellets, which was independent of intestinal motility (Figures 4B and S6). Treatment of mutant mice with vehicle, DPP4 inhibitor diluted in water, prolonged their survival but did not impact their fecal output or basal electrophysiology (Figure 4A-C), consistent with therapeutic administration of supportive fluids in diarrheal disease.

**Figure 4.**
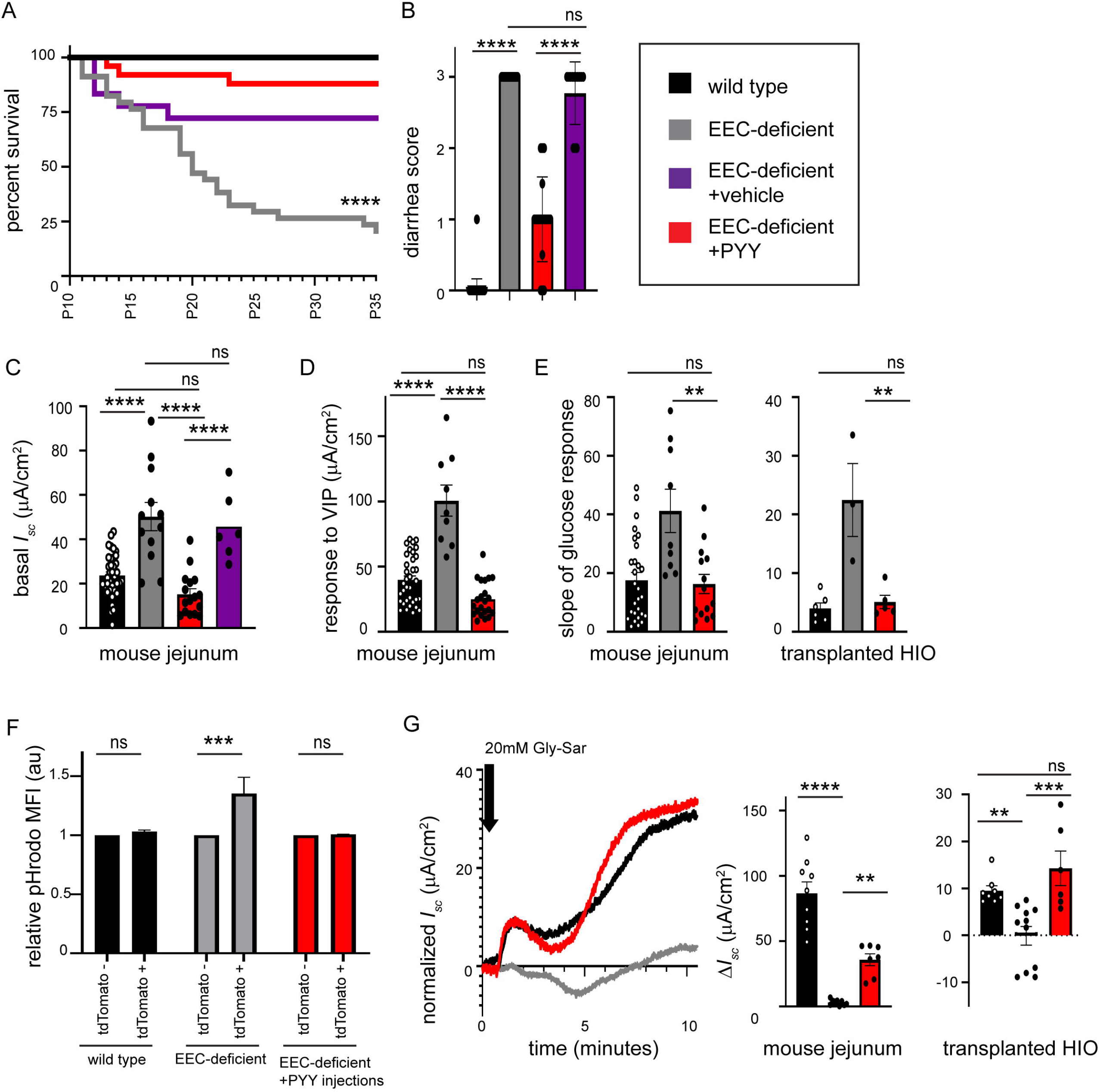
Exogenous PYY rescues EEC-deficient mice from malabsorptive diarrhea and death and restores normal glucose and dipeptide transport. A. PYY treatment promotes survival of EEC-deficient mice. Survival curve of wild-type (n=100), EEC-deficient (n=34) and EEC-deficient mice treated once daily with 10 μg PYY (n=25) beginning at postnatal day 10 (P10). Vehicle-treated mice received DPP4 inhibitor diluted in 100μl water (n=18). Mice were weaned at P21. Statistics calculated by log-rank Mantel-Cox test. B. Daily treatment of EEC-deficient mice with PYY reverses intractable diarrhea. As compared to control, EEC-deficient mice have intractable watery diarrhea from birth (given score of 3, gray bar; n=34; ****P<0.0001). Within 48 hours of PYY treatment, EEC-deficient animals had an average score of 1 with slightly soft yet well-defined fecal pellets (n=25, ****P<0.0001 from untreated mutant). Mutant mice treated with vehicle did not gain improvement in diarrhea score (n=18, ****P<0.0001 from PYY-treated mutant, and not significant from untreated mutant). Wild-type littermates produce well-defined fecal pellets (given score of 0, black bar; n=100). Error bars are ± SEM; statistics calculated by one-way ANOVA with Tukey’s multiple comparisons test. C. PYY treatment of EEC-deficient animals restored a normal resting *I*_sc_ to small intestine. Jejunum from wild-type (black), *VillinCre; Neurog3^flox/flox^* (gray), *VillinCre; Neurog3^flox/flox^* + PYY injected (red) and *VillinCre; Neurog3^flox/flox^* + vehicle injected (purple) mice were mounted in the Ussing chamber. Mutant jejunum exhibited a significantly increased basal *I*_sc_ compared to wild-type, which was significantly decreased after *in vivo* injections of PYY (n=6, ****P<0.0001). Treatment of mutant mice with vehicle did not result in improved basal *I*_sc_ (n=6). Wild-type and untreated mutant data points are the same as Supplemental Figure 2. Error bars are ± SEM; statistics calculated by one-way ANOVA with Tukey’s multiple comparisons test. D. Electrogenic response to VIP was elevated in EEC-deficient animals but restored to wild-type levels in mutant mice treated with PYY (n=6, ****P<0.0001). Wild-type and untreated mutant data points are the same as Figure 1. Error bars are ± SEM; statistics calculated by one-way ANOVA with Tukey’s multiple comparisons test. E. PYY treatment restores a normal glucose response in EEC-deficient mouse and human intestine. (mouse, n=6, **P=.003; HIO, n=5, **P=0.004). Wild-type and untreated mutant data points are the same as Figure 2. Error bars are ± SEM; statistics calculated by one-way ANOVA with Tukey’s multiple comparisons test. F. Proton transport is normalized in EEC-deficient animals following PYY treatment. Mean fluorescent intensity (MFI) of pHrodo was normalized between EEC-deficient and EEC-rich regions of the mosaic jejunum (n=2). MFI was normalized to untreated wild-type. Wild-type and untreated mutant data points are the same as Figure 3. Error bars are ± SEM; statistics calculated by two-way ANOVA with Sidak’s multiple comparisons test. G. PYY improves dipeptide transport in EEC-deficient mouse and human intestine. Long-term treatment of EEC-deficient animals and animals hosting transplanted HIOs with PYY resulted in improved *I*_sc_ response to luminal Gly-Sar compared to untreated mutant tissue (mouse, n=6, **P=.009; HIO, n=5, ***P=0.0001). Wild-type and untreated mutant data points are the same as Figure 3. Error bars are ± SEM; statistics calculated by one-way ANOVA with Tukey’s multiple comparisons test.

We investigated if the animals that survived in response to PYY injections had restored electrophysiology and improved nutrient absorption in the small intestine. We found that PYY-injections restored the basal *I*_sc_ of jejunum to normal (Figure 4C). Additionally, the response to VIP (Figure 4D) and the response to luminal glucose (Figure 4E) were both normalized indicating that PYY injections stably restored electrophysiology. Importantly, mice received their last injection of PYY approximately 16 hours prior to sacrifice, demonstrating sustained action of the peptide *in vivo*. The rescue of EEC-deficient intestinal tissue also extended to the human model, where EEC-deficient HIOs were grown and matured *in vivo* and then host animals were injected with exogenous PYY for 10 days prior to harvest. These EEC-deficient HIOs exposed to PYY demonstrated electrogenic response to glucose that was indistinguishable from wild-type (Figure 4E). Lastly, we investigated whether the PYY treated groups had improved amino acid absorption as measured by H^+^ export and response to the dipeptide Gly-Sar. By administering PYY to the mosaic EEC-deficient reporter mice, we found PYY injections restored intracellular pH in EEC-deficient intestinal cells to normal levels which would support PEPT1-mediated dipeptide absorption (Figure 4F). Consistent with this, PYY-injected mouse and human small intestine displayed a significantly improved electrogenic response to dipeptides (Figure 4G), indicating that dipeptide absorption was restored. These data demonstrated functional efficacy of PYY on improved ion and nutrient transport in EEC-deficient intestine.

## Discussion

In this study, we found that loss of all EECs resulted in profound imbalance of ion transport in the small intestine, with subsequent impairment of nutrient absorption. We demonstrated that PYY is an essential regulator of normal electrophysiology and absorptive function in the small intestine. Chemical inhibition of the epithelial NPY1R receptor in wild-type small intestine isolated from human intestinal organoids and mouse demonstrated the requirement of this pathway in the modulation of VIP-induced ion secretion. Administration of PYY to EEC-deficient animals resulted in improvements in survival, diarrheal symptoms, glucose absorption and protein absorption in the absence of all other EEC peptides.

Historically, mouse models have been exceedingly tolerant of loss of individual EEC populations, largely due to functional overlap between EEC-derived peptides (McCauley, 2019). This has rendered it difficult to assign roles of individual EEC peptides to physiologic functions. Here, we were able to exploit a model which lacks all EECs to functionally evaluate the role of one EEC peptide, PYY. However, other peptides like somatostatin have similar activities to PYY and likely play a similar regulatory role *in vivo*. Somatostatin has many systemic targets (Patel, 1999) and the use of the somatostatin-analogue octreotide in the treatment of chylous effusion and hyperinsulinemia causes an increased risk of necrotizing enterocolitis in infants (Chandran et al., 2020). We therefore chose to use PYY in our preclinical model of malabsorptive diarrhea.

PYY has been classically defined as a satiety hormone that acts in an endocrine manner wherein the DPP4-cleaved PYY(3-36) signals to the brain to reduce food intake (Batterham et al.). However PYY(1-36) has been shown to act in a paracrine manner in the colon using combination of genetic and pharmacological approaches (Cox, 2008; Hyland et al., 2003; Tough et al., 2011). We and others (Egerod et al., 2012) observe abundant PYY+ cells in the small bowel, suggesting that these cells may reprise their paracrine role described in the colon in the regulation of ion and water transport in the small intestine, linking EECs to glucose and protein absorption. These findings lend some clarity on how EECs integrate their nutrient sensing function with nutrient absorption, providing us with a new way to approach management of malabsorptive diseases and those in which EECs are commonly dysregulated.

## Figure Legends

**Supplemental Figure 1.**
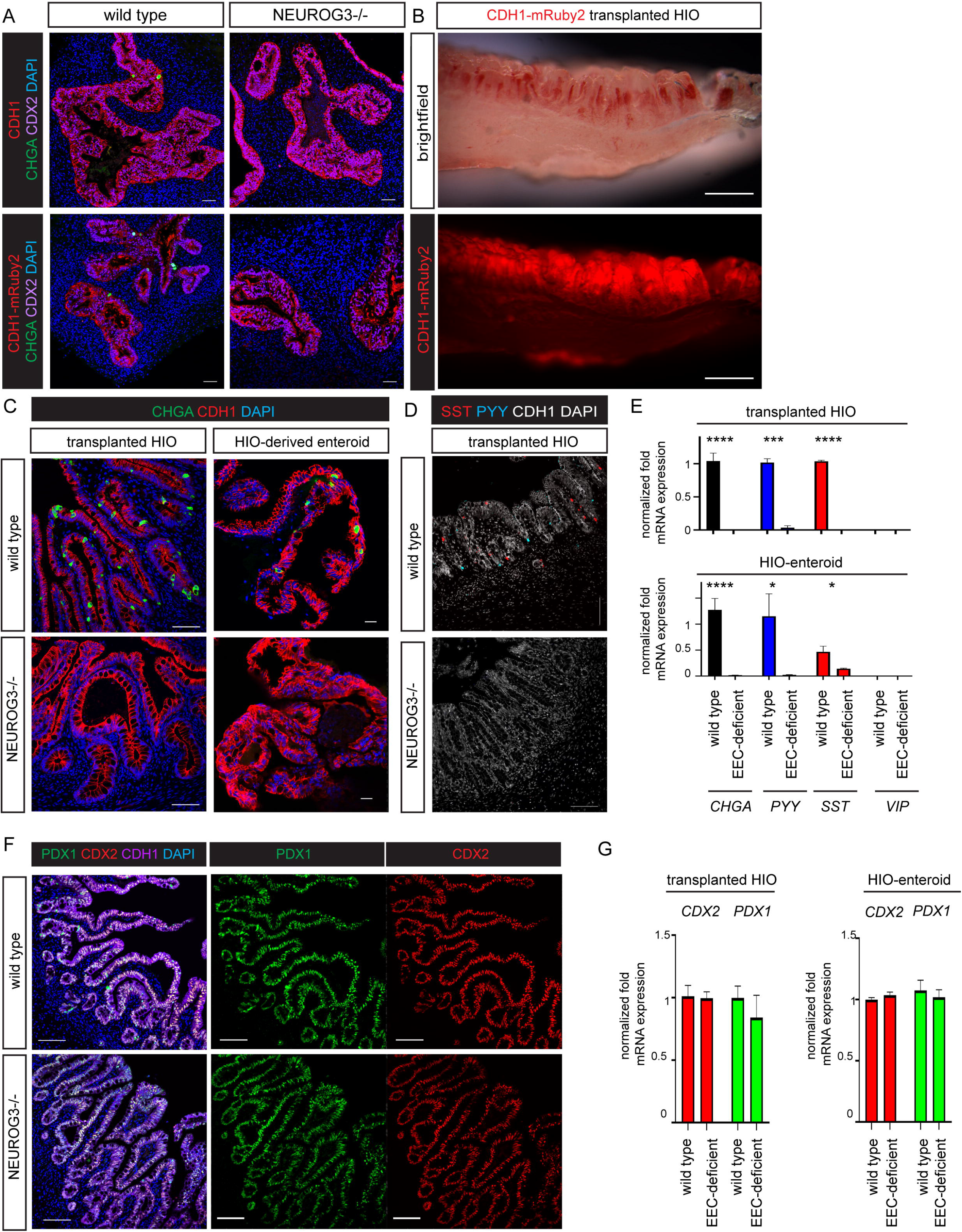
NEUROG3 is required for enteroendocrine cell development in human intestinal organoids. A. Human intestinal organoids (HIOs) derived from human pluripotent stem cells with a null mutation in *NEUROG3* lacked enteroendocrine cells (EECs) but otherwise had a normal morphology. The epithelial morphology was assessed using a PSC line expressing a CDH1-mRuby2 fusion protein (Ouchi et al., 2019) (red, bottom panels) and by co-staining with an anti-CDH1 antibody (red, top panels). Loss of NEUROG3 did not alter markers of intestinal identity (CDX2, purple). Only wild-type (top) and wild-type CDH1-mRuby2 (bottom) HIOs generated Chromogranin A (CHGA)- expressing EECs (green). Scale bars = 50 μm. B. After maturation *in vivo*, HIOs develop well-defined crypt-villus architecture. Transplantation of HIOs (~1 mm) into mice for 10-12 weeks results in growth (1-2 cm), morphogenesis and maturation (Watson et al., 2014). The epithelium is labeled by CDH1-mRuby2. Scale bar = 500 μm. C. Transplanted HIOs with disrupted *NEUROG3* lacked EECs as marked by CHGA+, but were otherwise morphologically normal. Scale bars = 100 μm. Enteroids derived from the crypts of transplanted HIOs produced EECs when differentiated, whereas those derived from EEC-deficient HIOs never did. DAPI and CDH1 mark nuclei and epithelium, respectively. Scale bars = 20 μm. D. Transplanted HIOs generated PYY+ and somatostatin (SST)+ EECs, which were never detected in NEUROG3-deficient transplanted HIOs. DAPI and CDH1 mark nuclei and epithelium, respectively. Scale bars = 100 μm. E. EEC-deficient transplanted HIOs (top) and EEC-deficient HIO-derived enteroids (bottom) did not express mRNA for EEC markers *CHGA* (****P<0.001), *PYY* (***P=0.001) or *SST* (****P<0.0001). Neither wild-type nor EEC-deficient tissues expressed mRNA for *VIP* (n=9). Error bars are ± SEM.; statistics calculated by unpaired, two-tailed Student’s *t*-test. F. Regional patterning of transplanted HIOs was independent of NEUROG3. Transplanted HIOs, with and without EECs, coexpressed CDX2 and the proximal small intestinal marker PDX1. DAPI and CDH1 mark nuclei and epithelium, respectively. Scale bars = 100 μm. G. Regional identity of transplanted HIOs was maintained in enteroid culture. There was no difference in *CDX2* or *PDX1* mRNA expression between wild-type and EEC-deficient transplanted HIOs, or between wild-type and EEC-deficient HIO-derived enteroids. Error bars are ± SEM.; statistics calculated by unpaired, two-tailed Student’s *t*-test.

**Supplemental Figure 2.**
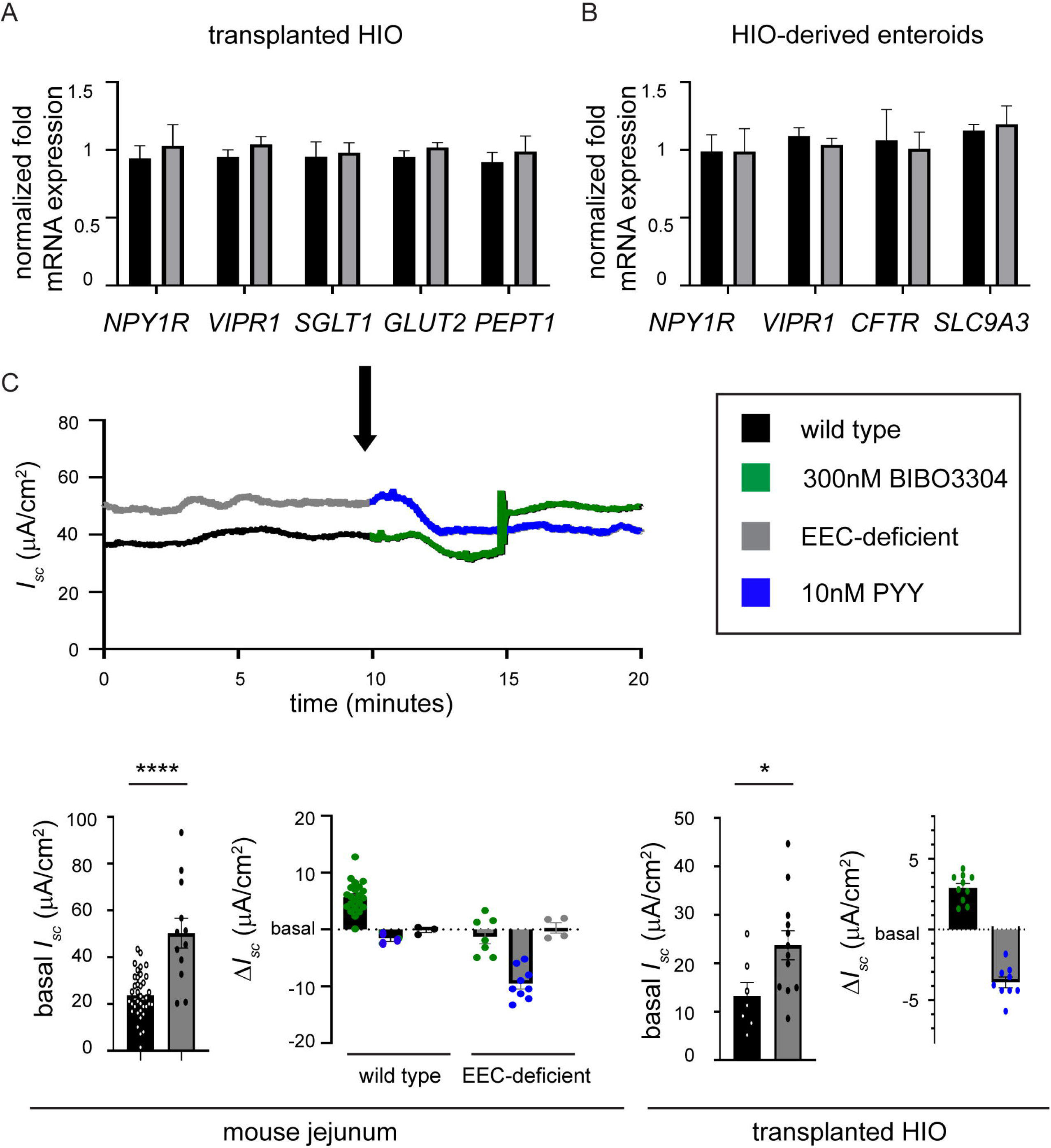
PYY is required to maintain normal electrophysiology in mouse and human small intestine. A. There was no difference in *NPY1R, VIPR1, SGLT1, GLUT2,* or *PEPT1* mRNA expression between transplanted HIOs with EECs and those without EECs. Error bars are ± SEM. B. There was no difference in *NPY1R, VIPR1, CFTR* or *SLC9A3* mRNA expression between enteroids generated from wild-type or EEC-deficient HIOs. Error bars are ± SEM. C. PYY modulates basal *I*_sc_ in human and mouse small intestine. EEC-deficient mouse and human small intestine had significantly higher basal *I*_sc_ than wild-type (mouse, n=36 wild-type, n=11 mutant, ****P<0.0001; HIO, n=7 wild-type, n=12 mutant, *p=0.03) after equilibration in the Ussing chamber. Addition of 300 nM NPY1R inhibitor BIBO3304 to wild-type tissues reproducibly increased the basal *I*_sc_ (mouse, n= 26, human, n=10), whereas addition of 10 nM PYY lowered the basal *I*_sc_ in mutant mouse and human tissue (mouse, n=9, human, n=9). Blocking NPY1R with BIBO3304 abolished the effect of PYY in both wild-type and mutant tissues. Arrow indicates time of PYY or BIBO3304 application to the experiment. One representative trace is shown (mouse). Error bars are ± SEM; statistics calculated by unpaired, two-tailed Student’s *t*-test.

**Supplemental Figure 3.**
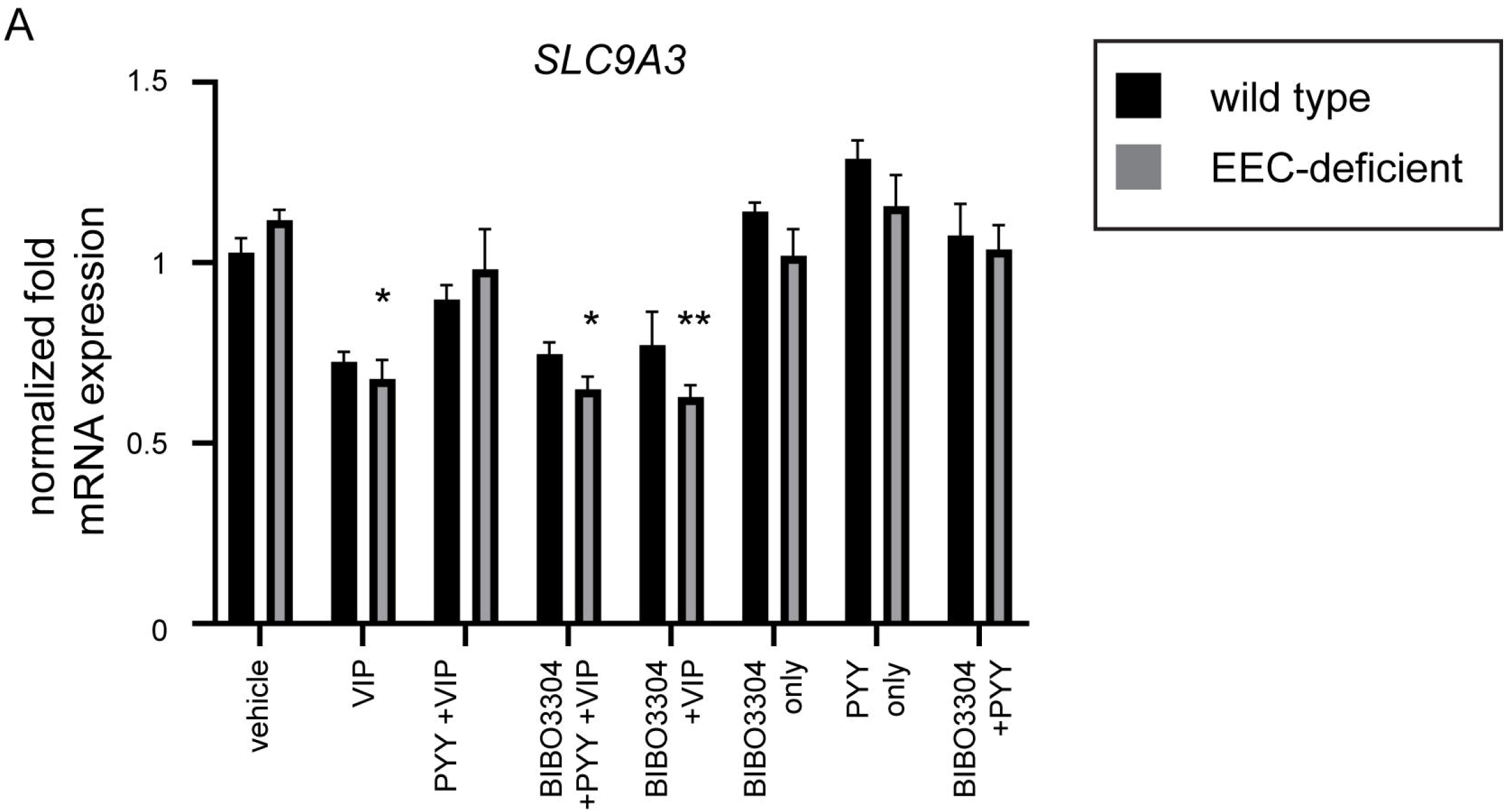
VIP and PYY regulate NHE3 expression in human small intestinal epithelium. A. The PYY-VIP axis regulates *SLC9A3* expression. After 5-7 days of exposure to VIP, *SLC9A3* expression was reduced in wild-type and in EEC-deficient enteroids (*P=0.04). Exposure to PYY concurrently with VIP in EEC-deficient enteroids restored *SLC9A3* expression to not significantly different from untreated. The effect of PYY was blocked with the NPY1R inhibitor BIBO3304 (*P=0.02). While there was a trend for PYY treatment alone to increase *SLC9A3* expression, this did not reach significance. n=6 independent experiments. Error bars are ± SEM; statistics calculated by one way ANOVA with Tukey’s multiple comparisons test.

**Supplemental Figure 4.**
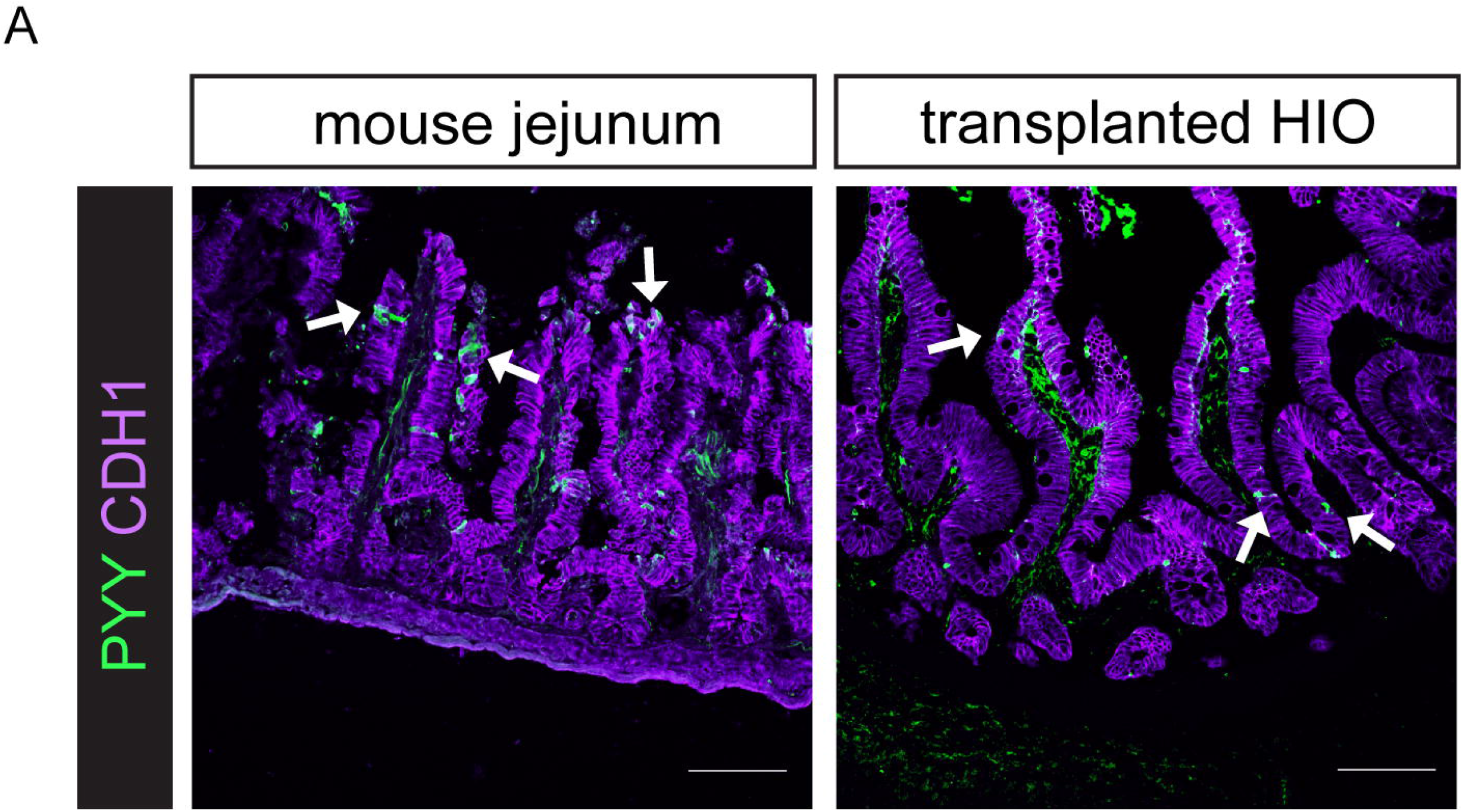
PYY is abundant in mouse and human small intestine. A. PYY+ EECs (arrows) are abundant in mouse and human small intestine. CDH1 labels epithelium in purple. Scale bars = 100 μM.

**Supplemental Figure 5.**
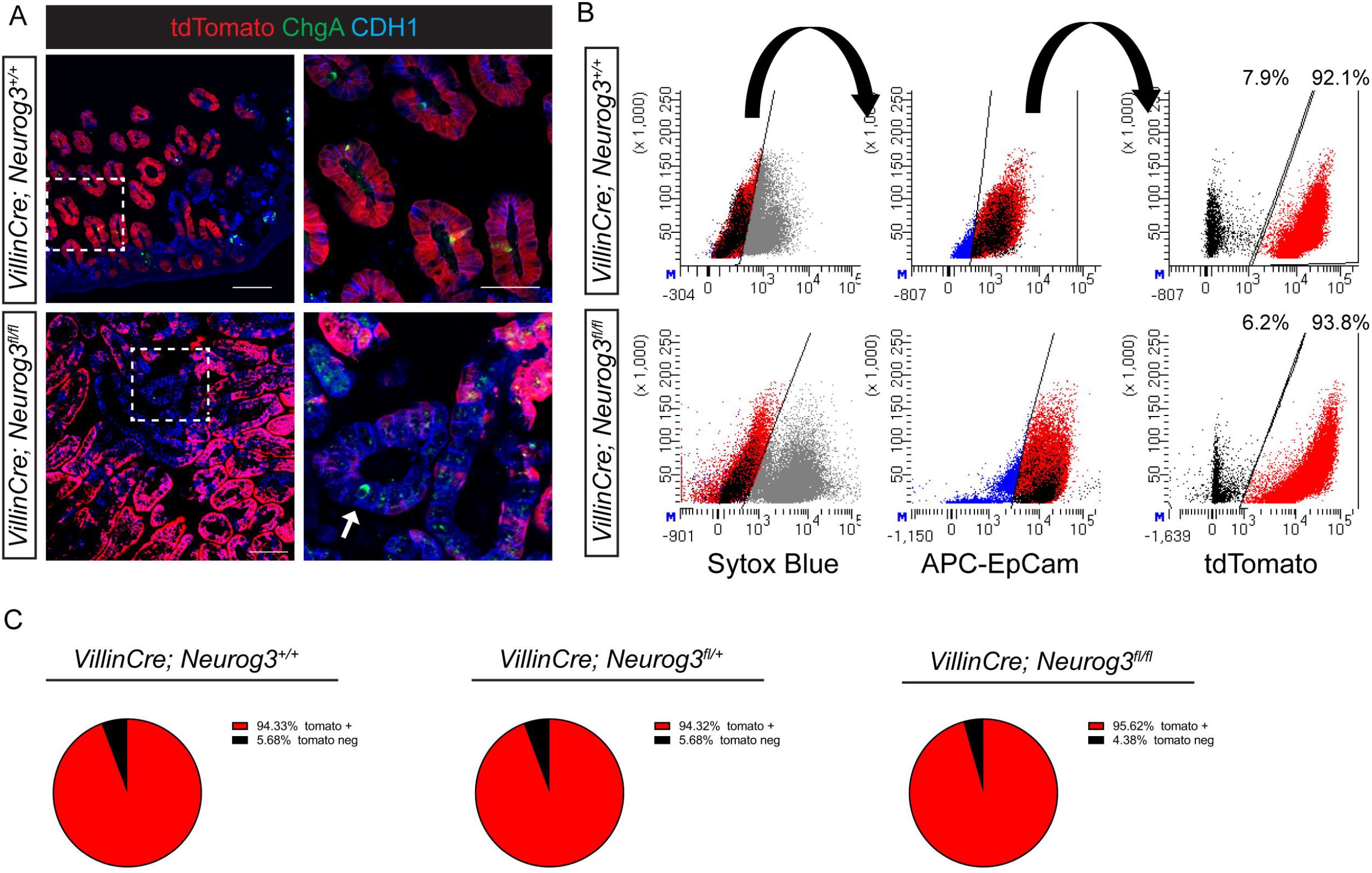
*VillinCre; Neurog3^flox/flox^; Rosa26^Flox-STOP-flox-tdTomato^* mice display incomplete recombination. A. The tdTomato reporter revealed regions of jejunal epithelium that escaped recombination by *VillinCre.* ChgA+ EECs were abundant in tdTomato+ and tdTomato negative regions of wild-type jejunum, but were only detected in tdTomato negative epithelium of *Neurog3^fl/fl^* animals (arrow). Scale bars = 20μm. B. Representative dot plots and gating strategy from flow cytometric analysis of *VillinCre; Neurog3^+/+^*; *Rosa26^Flox-STOP-flox-tdTomato^* and *VillinCre*; *Neurog3^flox/flox^*; *Rosa26^Flox-STOP-flox-tdTomato^* jejunum. C. Quantification of efficiency of recombination of *VillinCre*. Jejunum of *VillinCre*; *Neurog3^+/+^*; *Rosa26^Flox-STOP-flox-tdTomato^*, *VillinCre*; *Neurog3^fl/+^*; *Rosa26^Flox-STOP-flox-tdTomato^* and *VillinCre*; *Neurog3^fl/fl^*; *Rosa26^Flox-STOP-flox-tdTomato^* were subjected to flow cytometry. After doublet discrimination, live, EpCam^+^ cells were analyzed for tdTomato expression. Approximately 5.675 ± 1.98% of wild-type (n=8), 5.678 ± 3.2% of heterozygous (n=9), and 4.38 ± 2.56% of mutant jejunum (n=5) escaped labeling with the tdTomato reporter.

**Supplemental Figure 6.**
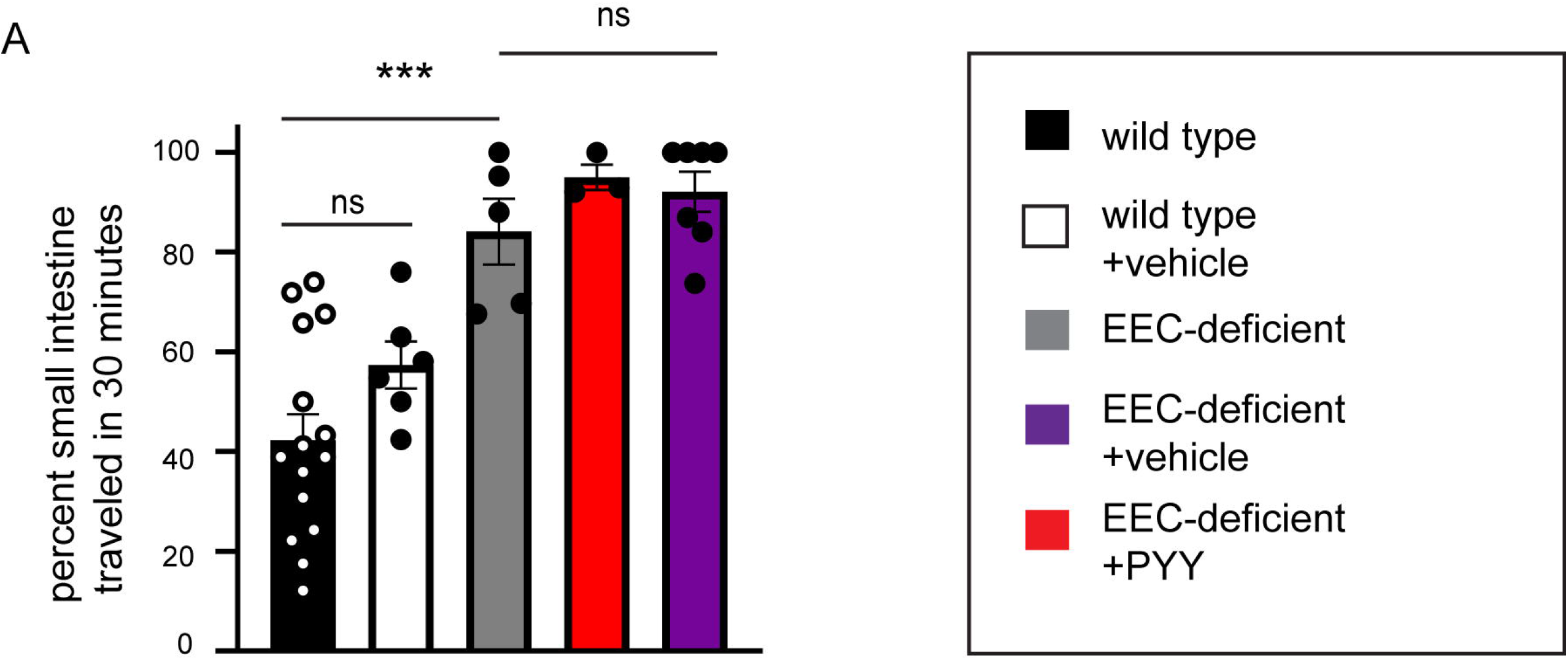
PYY(1-36) does not slow intestinal motility in EEC-deficient mice. A. The mechanism of improved survival and diarrhea in PYY-treated mutant animals does not include slowing intestinal motility. Animals fed ad-lib were orally gavaged with dye-colored water then sacrificed 30 minutes later. The distance traveled by the dye-front was reported as percent of small intestinal length. n=15 wild-type mice, 6 wild-type + vehicle mice, 5 mutant mice (***P=0.0002), 3 mutant + PYY mice, 7 mutant + vehicle mice. Error bars are ±SEM. Statistics calculated by one-way ANOVA with Tukey’s multiple comparisons test.

## Methods

### Pluripotent stem cell culture and directed differentiation of HIOs

Human embryonic stem cell (ESC) line WA01 (H1) was purchased from WiCell. We used H1 cells with a CRISPR/Cas9 generated null mutation in *NEUROG3* as previously described (McGrath et al., 2015). Additionally, we inserted the CDH1-mRuby2 reporter construct (Ouchi et al., 2019) into *NEUROG3−/−* H1 hESCs. CDH1-mRuby2 and non-reporter hESCs were used interchangeably. hESCs were maintained in feeder-free culture. Cells were plated on hESC-qualified Matrigel (BD Biosciences, San Jose, CA) and maintained at 37 °C with 5% CO_2_ with daily removal of differentiated cells and replacement of mTeSR1 media (STEMCELL Technologies, Vancouver, Canada). Cells were passaged routinely every 4 days using Dispase (STEMCELL Technologies). HIOs were generated according to protocols established in our lab (Múnera and Wells, 2017; Spence et al., 2011).

### *In vivo* transplant of HIOs

28-35 days after spheroid generation, HIOs were removed from Matrigel and transplanted under the kidney capsule of immune deficient NOD.Cg-*Prkdc^scid^Il2rg^tm1Wjl^*/SzJ (NSG) mice as previously described (Watson et al., 2014). NSG mice were maintained on Bactrim chow for a minimum of 2 weeks prior to transplantation and thereafter for the duration of the experiment (8-14 weeks).

### Generation and maintenance of HIO-derived enteroids

After approximately 10 weeks of *in vivo* growth, crypts were isolated from transplanted HIOs and plated in 3D as previous described (Mahe et al., 2015). To promote growth, enteroids were maintained in Human IntestiCult components A+B (STEMCELL Technologies). To promote differentiation, HIOEs were cultured in gut media (Múnera and Wells, 2017) with 100 μg/ml EGF for 5-7 days. Undifferentiated enteroids were passaged every 7-10 days into fresh Matrigel (Corning) using a 25G x1/2 needle.

### Immunofluorescence

Tissue was fixed in 4% paraformaldehyde, cryopreserved in 30% sucrose, embedded in OCT, and frozen at −80 °C until cryosectioned. 8 μm cryosections were mounted on Superfrost Plus slides and permeabilized, blocked and stained according to standard protocol. Primary antibodies used are listed in the table below, and all secondary antibodies were conjugated to Alexa Fluor 488, 546/555/568 or 647 (Invitrogen). Images were acquired using a Nikon A1 GaAsP LUNV inverted confocal microscope and NIS Elements software (Nikon).

| *Primary antibody* | *Company* | *Host* | *Dilution* |
| --- | --- | --- | --- |
| CDX2 | BioGenex | Mouse | 1:300 |
| CDX2 | Cell Marquis | Rabbit | 1:500 |
| Chromogranin A | DSHB | Mouse | 1:500 |
| Chromogranin A | ImmunoStar | Rabbit | 1:250 |
| E-Cadherin (CDH1) | R&D | Goat | 1:500 |
| GLUT2 | Santa Cruz | Goat | 1:500 |
| Muc2 | Santa Cruz | Rabbit | 1:250 |
| NPY1R | Abcam | Rabbit | 1:250 |
| PDX1 | Abcam | goat | 1:5000 |
| PEPT1 | Santa Cruz | Rabbit | 1:500 |
| PYY | Abcam | Rabbit | 1:1000 |
| SGLT1 | Santa Cruz | Rabbit | 1:250 |
| Somatostatin | Santa Cruz | Goat | 1:200 |
| VIPR1 | Millipore | Mouse | 1:200 |

### qPCR

RNA was extracted using Nucleospin RNA extraction kit (Macharey-Nagel) and reverse transcribed into cDNA using Superscript VILO (Invitrogen) according to manufacturer’s instruction. qPCR primers were designed using NCBI PrimerBlast. Primer sequences are listed in the table below. qPCR was performed using Quantitect SYBR® Green PCR kit (QIAGEN) and a QuantStudio 3 Flex Real-Time PCR System (Applied Biosystems). Relative expression was determined using the ΔΔCt method and normalizing to PPIA (cyclophilin A). Samples from at least three independent passages were used for quantification.

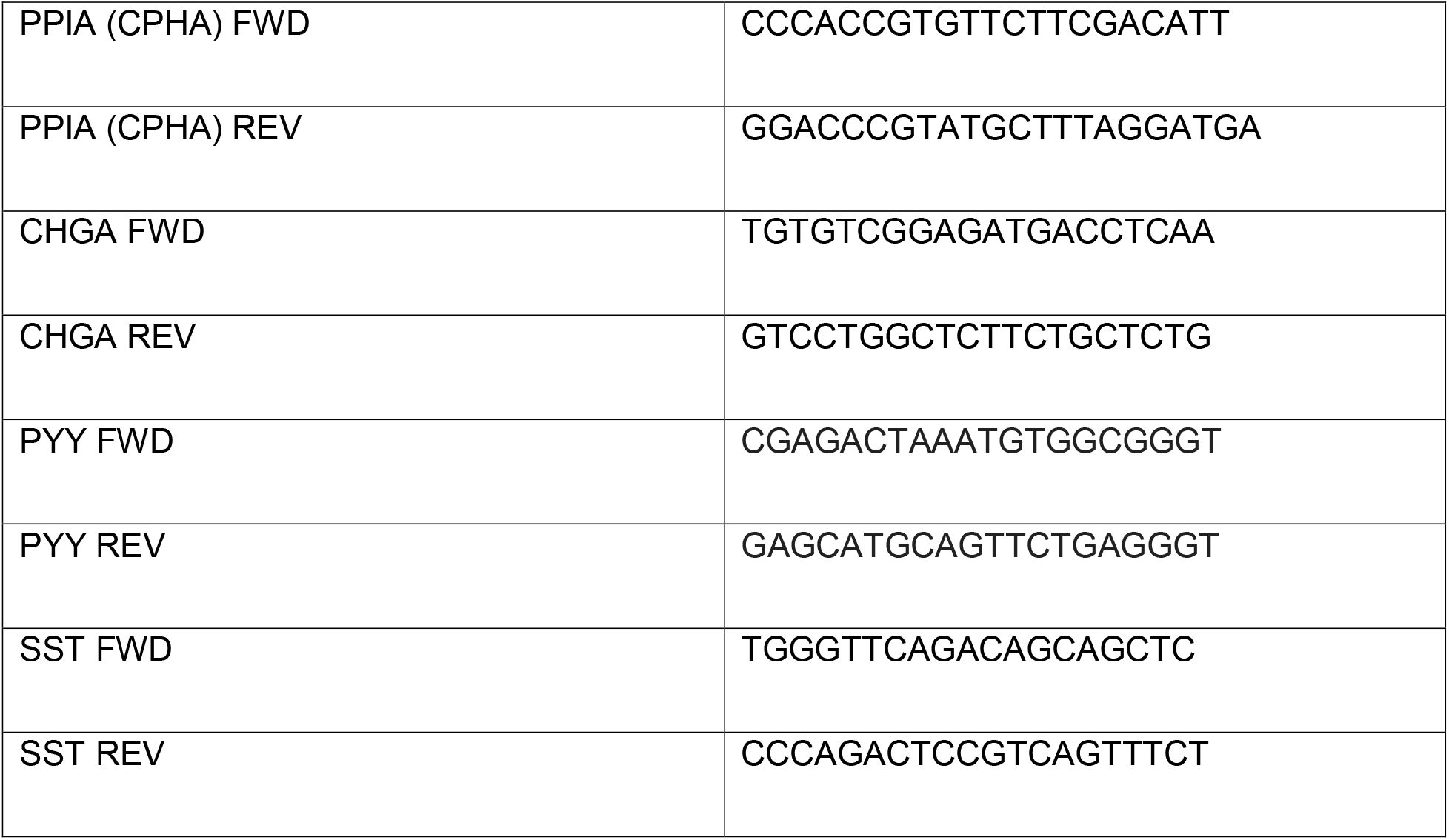
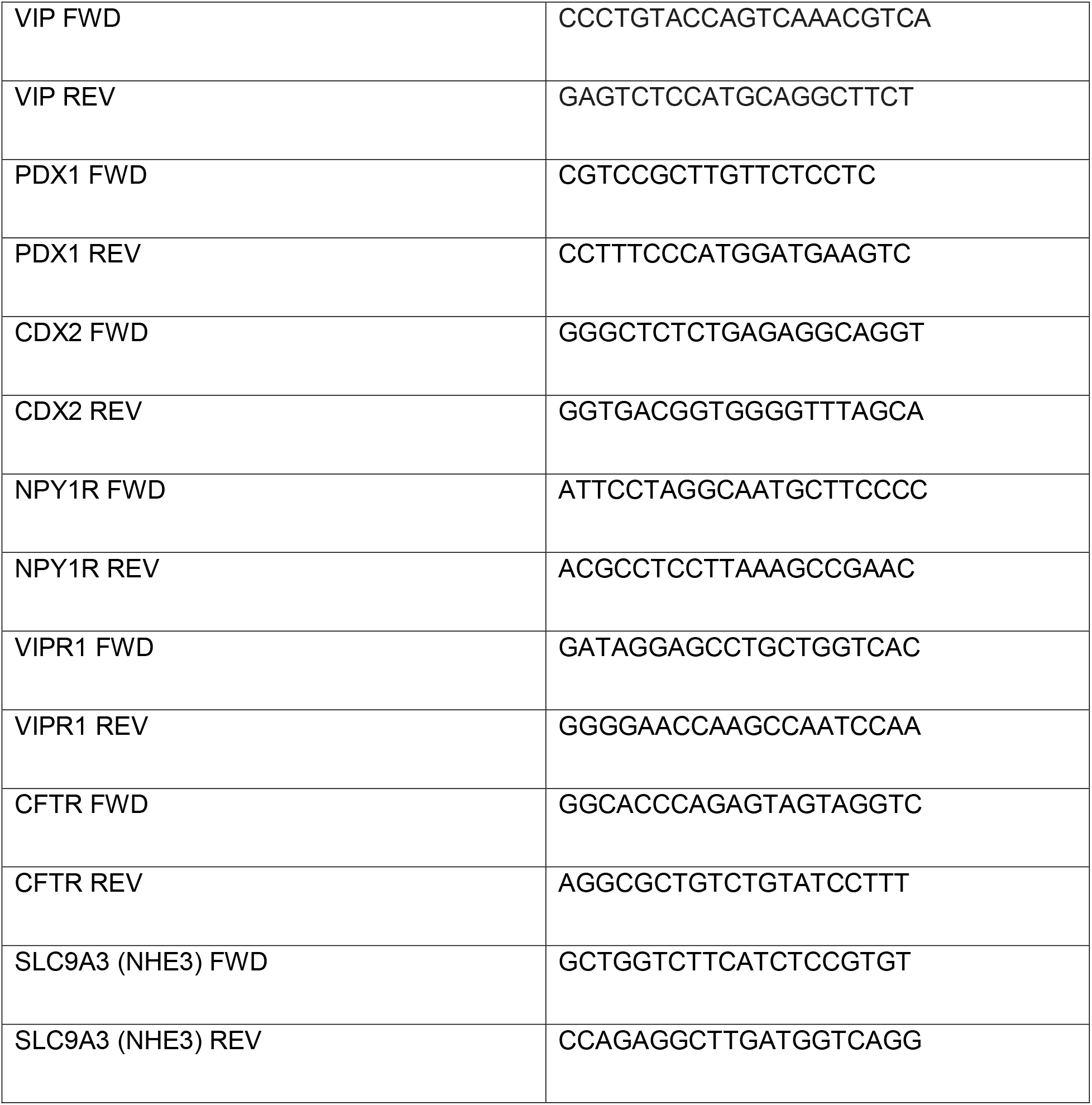

### Swelling assay

Enteroids were plated in 10 μL Matrigel on an 8-chamber glass bottom slide (Ibidi) and maintained as described above. 3-5 days post-plating, the slide was mounted on an inverted confocal microscope (Nikon) fitted with an incubation chamber set to 37 °C and 5% CO_2_. Media was changed to include 10 nM VIP (Tocris). In some experiments, the media was changed 24 hours prior to imaging to include 300nM BIBO3304 trifluoroacetate (Tocris), 20 μM CFTR(inh)-172 (Millipore Sigma) and/or 10 nM PYY (Phoenix Pharmaceuticals). Images were acquired every 5 minutes at 4X magnification. After 6 hours, some HIOEs swelled to the point of bursting; therefore, we used images acquired at time 0 and at 6 hours for quantification. The area of 10 representative enteroids per well was quantified using NIS Elements software at both time points. The outline of individual enteroids was traced manually and the area calculated by NIS Elements. Fold change at 6 hours over baseline was reported. Data include a minimum of three independent experiments per condition on three wild-type and three EEC-deficient HIO-derived enteroid lines.

### NHE3 activity assay

NHE3 activity was determined as previously described (Foulke-Abel et al., 2016) with minor modifications. Enteroids were plated in 5 μL Matrigel on an 8-chamber glass bottom slide (Ibidi) and maintained as described above. 3-5 days post-plating, media was changed to Na^+^ media containing 5 μM SNARF-4F 5-(and-6)- carboxylic acid, acetoxymethyl ester, acetate (Molecular Probes) and allowed to incubate for 30 minutes. The slide was then mounted on an inverted confocal microscope (Nikon), fitted with an incubation chamber set to 37 °C and 5% CO_2_. Fresh Na^+^ media was provided before image acquisition. Images were acquired every 2 minutes for 2 hours at 10X magnification with excitation at 488 nm and emission at 561 nm and 640 nm. Media was changed to NH_4_Cl to acid-load the epithelium, then to tetramethylammonium (TMA) media to withdraw Na^+^. Na^+^ containing media was then added and NHE3 activity quantified as a measure of initial pH recovery. 1 mM probenecid and 5 μM SNARF were present in all buffers, and all buffers were set to pH 7.4. Intracellular pH was calibrated using the Intracellular pH Calibration Buffer kit (Invitrogen) at pH 7.5, 6.5 and 5.5 in the presence of 10 μM valinomycin and 10 μM nigericin at the conclusion of each experiment. The ratio of 561/640 was determined using NIS Elements software by drawing a region of interest and quantifying the fluorescence intensity of each wavelength over the period of the experiment. A minimum of 3 enteroids in 3 wells over two independent passages were quantified. The ratio of 561/640 was converted to intracellular pH using the equation provided by the manufacturer.

Na^+^ media: 130 mM NaCl, 5 mM KCl, 2 mM CaCl_2_, 1 mM MgSO_4_, 20 mM HEPES, 5 mM NaOH, 1 mM (Na)PO_4_, 25 mM D-glucose

NH_4_Cl media: 25 mM NH_4_Cl, 105 mM NaCl, 2 mM CaCl_2_, 1 mM MgSO_4_, 20 mM HEPES, 8 mM NaOH, 5 mM KCl, 1 mM (Na)PO_4_, 25 mM D-glucose

TMA media: 130 mM TMA-Cl, 5 mM KCl, 2 mM CaCl_2_, 1 mM MgSO_4_, 20 mM HEPES, 8 mM TMA-OH, 1 mM (TMA)PO_4_, 25 mM D-glucose

### Electrophysiology

Electrophysiological experiments were conducted as described (Clarke, 2009) with minor modifications. Mouse jejunum and transplanted HIOs were dissected and immediately placed in ice-cold Krebs-Ringer solution. Tissues were opened to create a flat epithelial surface. Because seromuscular stripping is associated with release of cyclooxygenases and prostaglandins (Clarke, 2009), and prostaglandins can stimulate L-cells to release GLP1, GLP2 and PYY (Briere et al., 2013), we performed the Ussing chamber experiments in intestinal tissue with an intact muscular layer. Tissues were mounted into sliders (0.031 cm^2^ area slider, P2307, Physiological Instruments) and placed in an Ussing chamber with reservoirs containing 5 mL buffer (115 mM NaCl, 1.2 mM CaCl_2_, 1.2 mM MgCl_2_, 25 mM NaHCO_3,_ 2.4 mM K_2_HPO_4_ and 0.4 mM KH_2_PO_4)_. The mucosal and serosal tissue surfaces were bathed in the same solution, with the exception of 10 mM glucose in the serosal buffer and 10 mM mannitol in the luminal buffer. Mucosal and serosal reservoir solutions were gassed with 95 % O_2_ and 5 % CO_2_ to pH 7.4 and maintained at 37 °C by a circulating water bath behind the reservoir chambers. Electrophysiology parameters were recorded as previously described (Matthis et al., 2019). Tissue was allowed to equilibrate to a basal steady-state for a minimum of 30 minutes before the addition of chemicals or peptides. 10 nM tetrodotoxin (Tocris) was added to the serosal buffer bathing mouse intestine to inhibit voltage-gated neuronal firing, and allowed to incubate for a minimum of 10 minutes before basal *I*_sc_ recording. D-glucose and Gly-Sar were added to the luminal side of the chamber once the VIP-induced *I*_sc_ had stabilized at a maximum value.

|  |  |  |
| --- | --- | --- |
| Tetrodotoxin | Tocris | 10 nM |
| BIBO3304 trifluoroacetate | Tocris | 300 nM |
| VIP | Tocris | 10 nM |
| PYY(1-36) | Phoenix Pharmaceuticals | 10 nM |
| CFTR-172 | Millipore Sigma | 20 μM |
| D-glucose | Sigma Aldrich | 25 mM |
| Gly-Sar | Sigma Aldrich | 20 mM |

### Glucose uptake assays

#### 6-NBDG

Transplanted HIOs were removed from the murine kidney, bisected to expose the lumen, and incubated with 100 mM 6-(*N*-(7-Nitrobenz-2-oxa-1,3-diazol-4-yl)Amino)-2-Deoxyglucose (6-NBDG) (Life Technologies) in 10 nM Tris/HEPES buffer containing 150 mM KCl or 150 mM NaCl for 30 minutes at 37 °C. Tissues were washed with ice-cold 10 mM Tris/HEPES buffer, then dissociated to single-cell suspension in 5 mL Tryple Select (Gibco) + 10 μM Y-27632 (Tocris), filtered, and subjected to analysis by flow cytometry.

#### Sodium Green

HIOEs were differentiated for 5-7 days, then were removed from Matrigel and enzymatically dissociated into single-cell suspension using 0.25% Trypsin-EDTA. Each cell preparation was split into two samples: one incubated with 25 mM D-glucose and one incubated in the absence of glucose. Each sample was incubated in Live Cell Imaging Solution (Invitrogen) containing 5 μM final concentration of Sodium Green tetraacetate (Molecular Probes) for 30 minutes at 37 °C, washed with ice-cold PBS and analyzed by flow cytometry.

#### 2-NBDG on Transwell filters

Undifferentiated enteroids that were “ready to split” were dissociated and plated on transwell inserts (Corning) as previously described (Moon et al., 2014), with the exception of first coating the transwells with Collagen IV (Sigma-Aldrich). 300,000 cells were plated per 6.5 mm transwell insert. Differentiation was initiated at 24 hours post-plating and monolayers were analyzed after 5-7 days. 1 mM fluorescent glucose analog 2-(*N*-(7-Nitrobenz-2-oxa-1,3-diazol-4-yl)Amino)-2-Deoxyglucose (2-NBDG, Life Technologies) was diluted in Live Cell Imaging Solution (Invitrogen) containing 25 mM D-glucose, added to the apical surface of HIOE monolayers and the fluorescence intensity of fresh Live Cell Imaging Solution in the basal chamber was quantified after 30 minutes at 37 °C. Intact barrier function was confirmed by co-incubation, quantification and exclusion of Cascade Blue conjugated 3000 MW dextran (Life Technologies) in every experiment.

### Intracellular pH assay

Enteroids were differentiated for 5-7 days in the presence of vehicle (water or DMSO), 10 nM VIP (Tocris), 10 nM PYY (Phoenix Pharmaceuticals) and/or 300 nM BIBO3304. On the final day, enteroids were removed from Matrigel and enzymatically dissociated into single-cell suspension using 0.25% Trypsin-EDTA. Cell suspensions were counted and equal cell numbers of dissociated HIOEs were incubated in pHrodo Green AM Intracellular pH indicator (ThermoFisher Scientific) according to manufacturer’s directions for 30 minutes at 37C, washed with 1X PBS, and analyzed by flow cytometry.

### Flow cytometry

After mechanical and enzymatic dissociation, tissues were filtered through a 40 μm cell strainer to obtain a single-cell suspension. In all experiments, samples were labeled with either CDH1-mRuby2 or Anti-EpCam-APC (BD Biosciences) to distinguish epithelial cells and incubated with SYTOX Blue dead cell stain (Life Technologies) or 7-AAD (BD Pharmingen). Forward scatter and side scatter were used to discriminate doublets and cellular debris. A minimum of 50,000 events per sample was recorded using an LSR Fortessa flow cytometer (BD Biosciences) and data was analyzed using FACSDiva software (BD Biosciences).

### Mice

B6.Cg-*Tg(Vil1-cre)^997Gum/J^* (*VillinCre)* (JAX stock 004586)*, Neurog3^flox/flox^* (Mellitzer et al., 2010) and B6.Cg-*Gt(ROSA)26Sor^tm9(CAG-tdTomato)Hze^*/J (tdTomato) (Madisen et al., 2010) mice were maintained on a C57BL/6 background and genotyped as previously described. Mice were housed in a specific pathogen free barrier facility in accordance with NIH Guidelines for the Care and Use of Laboratory Animals. All experiments were approved by the Cincinnati Children’s Hospital Research Foundation Institutional Animal Care and Use Committee (IACUC2019-0006) and carried out using standard procedures. Mice were maintained on a 12-hour light/dark cycle and had *ad libitum* access to standard chow and water. *VillinCre;Neurog3^flox/flox^* mice^4^ and their littermates were weighed, genotyped and visually examined for liquid feces daily beginning at postnatal day 10. We established a diarrhea score, with 3 representing wet, yellow feces that smeared the perianal fur, and 0 representing normal, dry, brown, well-defined pellets. Mutant mice which suffered from diarrhea score 3 were included in the rescue experiment. 10 μg PYY (Phoenix Pharmaceuticals) was diluted in water and added to 20 μl DPP4 inhibitor (Millipore) to a final volume of 100 μl per mouse. Mice were injected intraperitoneally with this cocktail within 2 hours of the onset of the dark cycle (7pm) daily until analysis at postnatal day 28-35. Mice were given access to solid chow on the floor of the cage beginning at postnatal day 10 and weaned at postnatal day 21. Small intestinal transit was determined by oral gavage of food coloring diluted in 100μl to ad-lib fed mice, then sacrifice and measurement of the distance traveled by the dye-front 30 minutes post-gavage. NSG mice hosting HIOs were treated with 25 μg PYY (Phoenix Pharmaceuticals) diluted in water to 100 μL by intraperitoneal injection. Mice were treated daily for a minimum of 10 days after HIOs had been maturing for 8 weeks, then dissected and analyzed.

### Statistics

Data is presented as the mean ± SEM unless otherwise indicated. Significance was determined using appropriate tests in Graph Pad Prism, with P>0.05 not significant; *P<0.05, **P<0.01, ***P<0.001, ****P<0.0001.

## Acknowledgements

We thank Dr. Gerard Gradwohl and Dr. Andrew Leiter for providing the *Neurogenin3^flox/flox^* mice; Dr. Mary Estes, Dr. Sarah Blutt and Ms. Xi-Lei Zeng for training in generating HIO-derived enteroid monolayer culture systems; Ms. Catherine Martini for technical assistance. We acknowledge support provided by the Confocal Imaging Center, the Pluripotent Stem Cell Facility, and Research Flow Cytometry Core at CCHMC. We would like to thank the members of the Wells, Zorn, and Helmrath laboratories for reagents and feedback.

This work was supported by the grants from the NIH, U19 AI116491 (JMW), P01 HD093363 (JMW), UG3 DK119982 (JMW), U01 DK103117 (MAH); S&R Foundation and American Physiological Society (EA); the American Diabetes Association, 1-17-PDF-102 (HAM); and the Allen Foundation (JMW). We also received support from the Digestive Disease Research Center (P30 DK078392).

## Author contributions

HAM and JMW conceived and initiated the project, designed experiments, and wrote the manuscript, with conceptual input from MAH, MHM and EA. HAM performed all experiments in collaboration with: JRE, JGS and WJS on mouse transplantation; NS and MAH in generating HIO-derived enteroids; ALM, MHM and EA on electrophysiological studies. HAM, EA and JMW interpreted data. JMW supervised the project. All authors have edited and approved the manuscript.

## Materials and correspondence

Requests for materials and correspondence should be directed to.

